# Gut microbiome modulates behaviour and life history in two wild rodents

**DOI:** 10.1101/2020.02.09.940981

**Authors:** Joël W. Jameson, Denis Réale, Steven W. Kembel

**Author notes:** CORRESPONDING AUTHOR, Joël W. Jameson, Département des Sciences Biologiques, Université du Québec à Montréal, Montréal, Québec, Canada, H2X 1Y4.

## Abstract

Laboratory studies demonstrate that the gut microbiome can regulate host anxiety and exploratory behaviour. While this has implications for human health, it could have ecological and evolutionary implications for wild populations, a hitherto untested hypothesis. We tested whether the microbiome can directly modulate host behaviour and thereby affect life history in wild mice (*Peromyscus maniculatus*) and voles (*Myodes gapperi*). We compared the microbiome composition, exploration and anxiety behaviours, and home range of mice and voles before and after chronic antibiotic treatment and measured survival during treatment. Treated animals had lower diversity and relative abundance of most bacterial phyla save for Proteobacteria which increased in relative abundance. In mice, antibiotic treatment increased exploration and decreased home range without impacting survival. In voles, it lowered survival such that we could not test its effect on behaviour. Therefore, the microbiome can directly impact behaviour and host life history in the wild.

## INTRODUCTION

Modulation of host behaviour by the gut microbiome is well established from studies on laboratory mice and rats, relying on comparisons between germ free (GF) animals, antibiotic- or probiotic-treated animals, and animals with a conventional gut microbiome (Bercik et al., 2011; Desbonnet et al., 2014; Heijtz et al., 2011; Neufeld et al., 2011). Gut microbiome-mediated behavioural changes in these studies also correlate with neurochemical changes in the brain (Ait-Belgnaoui et al., 2012; De Palma et al., 2015; Desbonnet et al., 2014; Neufeld et al., 2011). While the gut microbiome has the potential to influence behaviour, this influence has only been studied in controlled laboratory experiments on genetically homogeneous groups of animals and in the context of studying human health and disease. However, the impact of the gut microbiome on behavioural traits such as anxiety and exploration could have important ecological and evolutionary consequences in wild populations.

In laboratory studies, GF mice show reduced anxiety-like behaviour (Heijtz et al., 2011; Neufeld et al., 2011), increased locomotion (Neufeld et al., 2011), increased social avoidance and reduced social cognition (avoidance of newly encountered conspecifics) (Desbonnet et al., 2014) compared with specific pathogen free (SPF) mice. These differences are accompanied by increased levels of plasma stress hormones and gene-level changes in the hippocampus and amygdala (Neufeld et al., 2011). In one study comparing GF and SPF mice, the microbiome was required for induction of an anxiety behaviour resulting from an early-life stressor (maternal separation) despite both groups presenting similar activity of the hypothalamic pituitary adrenal (HPA) axis (De Palma et al., 2015). The use of antibiotics has also allowed elucidation of important interactions between host behaviour and the microbiome in lab mice. Mice given non-absorbable antimicrobials have shown reduced anxiety and elevated exploration accompanied by elevated brain-derived neurotrophic factor (BDNF) in the hippocampus and reduced BDNF levels in the amygdala (Bercik et al., 2011). In a mouse model of anxiety, oral antibiotics suppressed the HPA stress response and promoted expression of proinflammatory cytokines in the hypothalamus (Ait-Belgnaoui et al., 2012). Finally, in a study by Gracias et al. (Gacias et al., 2016), offspring of C57BL/6J mice treated with antibiotics that lowered *Lactobacillus* and increased *Clostridium* abundances in the gut, exhibited reduced exploration and increased thigmotaxis in the open field (Tochitani et al., 2016). However, the same antibiotic treatment given to BALB/c mice for seven days increased exploratory behaviour, despite causing similar changes to the microbiome (Bercik et al., 2011), suggesting that the effect of the microbiome on behaviour can vary with species or even species strain.

In the wild, variation in anxiety and exploration-related behaviours can affect dispersal, reproduction, survival and fitness, with potential consequences for population structure (Ballew et al., 2017; Smith and Blumstein, 2008; Spiegel et al., 2017). Although these traits are often studied in the context of clinical behavioural disorders, variation in these traits also exists among individuals in wild populations (Réale et al., 2007) where they are often correlated, forming a behavioural syndrome (Réale et al., 2010). For example, animals with lower anxiety are more aggressive, less sociable, faster explorers (Aplin et al., 2013; Jolles et al., 2015; Kurvers et al., 2010; Réale et al., 2010), a pattern that is also observed when comparing GF mice to SPF mice (Desbonnet et al., 2014; Heijtz et al., 2011; Neufeld et al., 2011). Potential drivers of individual behavioural variation in the wild include genotype, resource competition, sexual selection, predation, and parasite infection (Barber and Dingemanse, 2010; Bell and Sih, 2007; Cote et al., 2008; Réale et al., 2007; Schuett et al., 2010). The gut microbiome may also play a key role in determining individual-level behavioural phenotype in the wild. Gut microbiome structure is correlated with level of social interaction in wild chimpanzees (Moeller et al., 2016) and social structure is also correlated with gut microbiome variation in wild baboons and lemurs (Diakiw, 2017; Tung et al., 2015). While this indicates a link between microbiome and social behaviour in the wild, we are not aware of any study that has empirically tested if the microbiome can modulate behaviour in wild animals. Given that the same behaviours under microbial control in the laboratory are also under selection in wild populations (Smith and Blumstein, 2008), the microbiome may also play a role in this latter context, though the extent of this role under natural ecological conditions remains to be determined.

We tested the hypothesis that the microbiome can directly modulate host exploration and anxiety-related behaviour in wild animals. We also aimed to determine whether a microbiome-induced change in host behaviour could influence aspects of host life history such as home range and survival. Finally, since behavioural response to microbial changes can vary with host species. We conducted this study on two sympatric rodent species, the deer mouse (*Peromyscus maniculatus*) and the red-backed vole (*Myodes gapperi*). Studying wild rodents, especially mice, allowed us to make more direct comparisons with results from laboratory studies. To address the goals of this study, we compared the microbiome, exploration and anxiety-related behaviours, and home range of wild mice and voles before and after chronic antibiotic treatment and measured their survival during the treatment period.

## RESULTS

We live-trapped mice and voles approximately every two days over a period of 41 days from 14 July – 23 August 2016. During a pre-treatment period (first 17 days) animals were captured but received no treatment. During the treatment period (day 18 to 41), animals were randomly assigned to either a group given saline, or a group given an antibiotic cocktail. We captured 19 male and 16 female deer mice, and 47 male and 28 female red-backed voles. Of the adult male mice, we gave a treatment of antibiotics to 10 and saline to eight. On average (± SD), mice received 5.7 ± 2.0 treatments and we treated mice every 2.6 ± 0.5 days. Of the adult male voles, 18 received antibiotics, and another 18 received saline. Voles received 3.9 ± 2.2 treatments and we treated them every 2.3 ± 0.7 days.

### Antibiotic Treatment Reduces Microbiome Diversity

We characterized the microbiome of mice and voles by sequencing the bacterial 16S barcode gene of two to three fecal samples per animal collected from traps throughout the study. We used DADA2 to identify amplicon sequence variants (ASV), our fundamental unit of ecological analysis (Callahan et al., 2016), and assigned taxonomy using the SILVA database (V128). We calculated Shannon’s diversity of each sample and Bray-Curtis dissimilarity among the samples (Oksanen et al., 2019). We then used principal coordinate analyses (PCoA) to plot pairwise dissimilarities in two dimensions and calculated the weighted average scores for taxa to identify those most associated with each PCoA axis (Oksanen et al., 2019).

The antibiotic treatment significantly changed the gut microbiome composition of both species (Fig. 1). In mice, variance along the first axis (PC1) was attributable to the antibiotic treatment, while variance along PC2 represented an unknown source, independent of the treatment (Figs. 2,3). In voles, the antibiotic treatment effect acted on both PC1 and PC2 (Figs. 2,3). Shannon’s diversity of both mice and voles decreased only in the antibiotic group from the pre-treatment to the treatment period, and the extent of this effect depended on the number of days since their last treatment (Fig. 3, Tables S1 and S2). PC1 increased from the pre-treatment to the treatment period only in the antibiotic-treated mice and voles, with this effect also varying as a function of the number of days since their last treatment (Fig. 3, Tables S1 and S2). Only saline-treated mice showed an increase along the PC2 axis from the pre-treatment to treatment period indicating that PC2 represented variation independent of the antibiotic treatment. In voles, variation along PC2 corresponded with the antibiotic treatment, increasing only in antibiotic-treated voles during the treatment period and once again, varying as a function of the number of days since the animal was last captured (Fig. 3, Tables S1 and S2).

**Fig.1.**
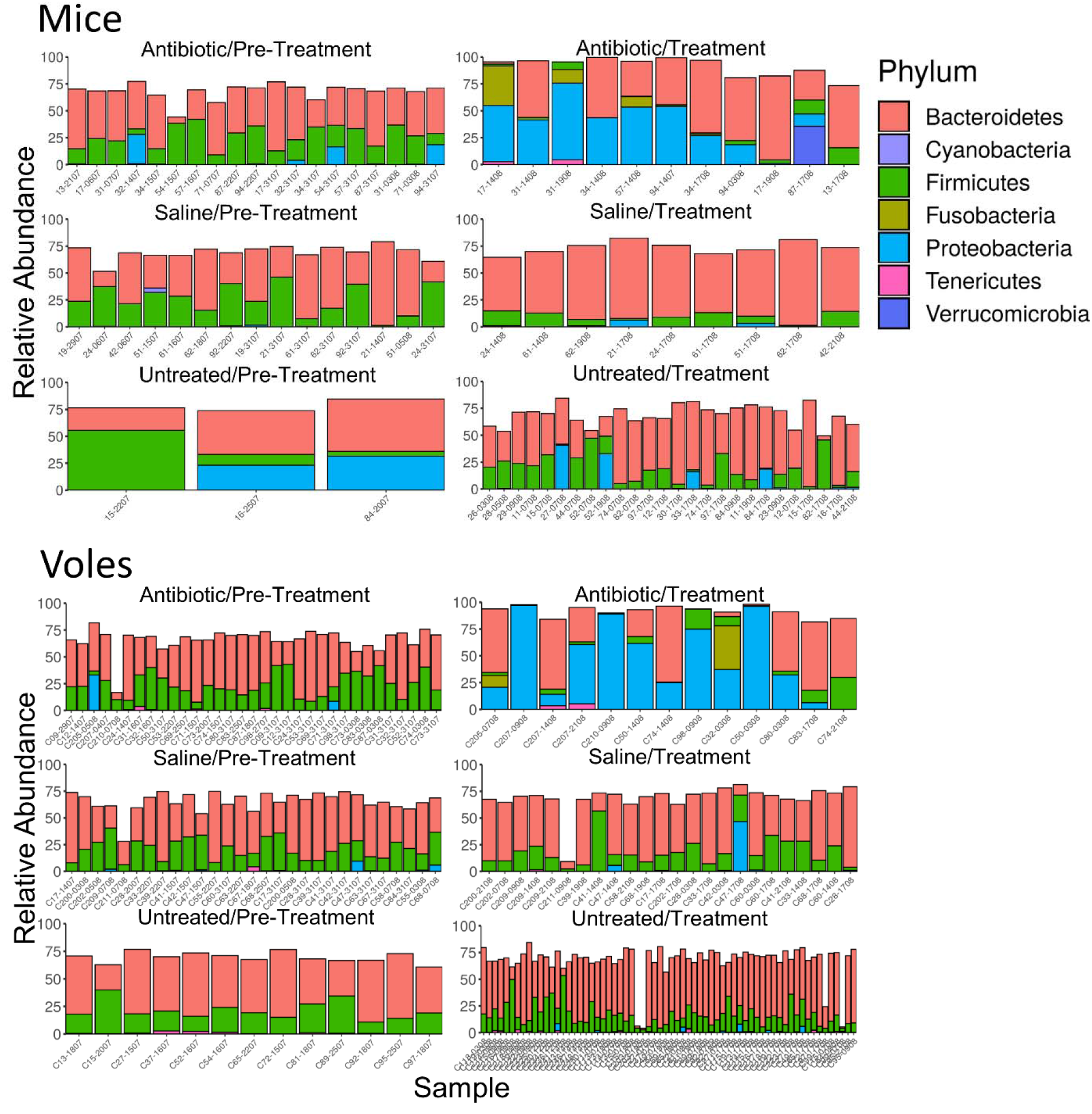
Effect of an antibiotic treatment on the composition of the gut microbiome in deer mice and red-backed voles. Relative abundance of the most abundant bacterial phyla in each sample. Samples are sorted by the number of days since the animal was last captured. Each bar represents an individual sample.

**Fig.2.**
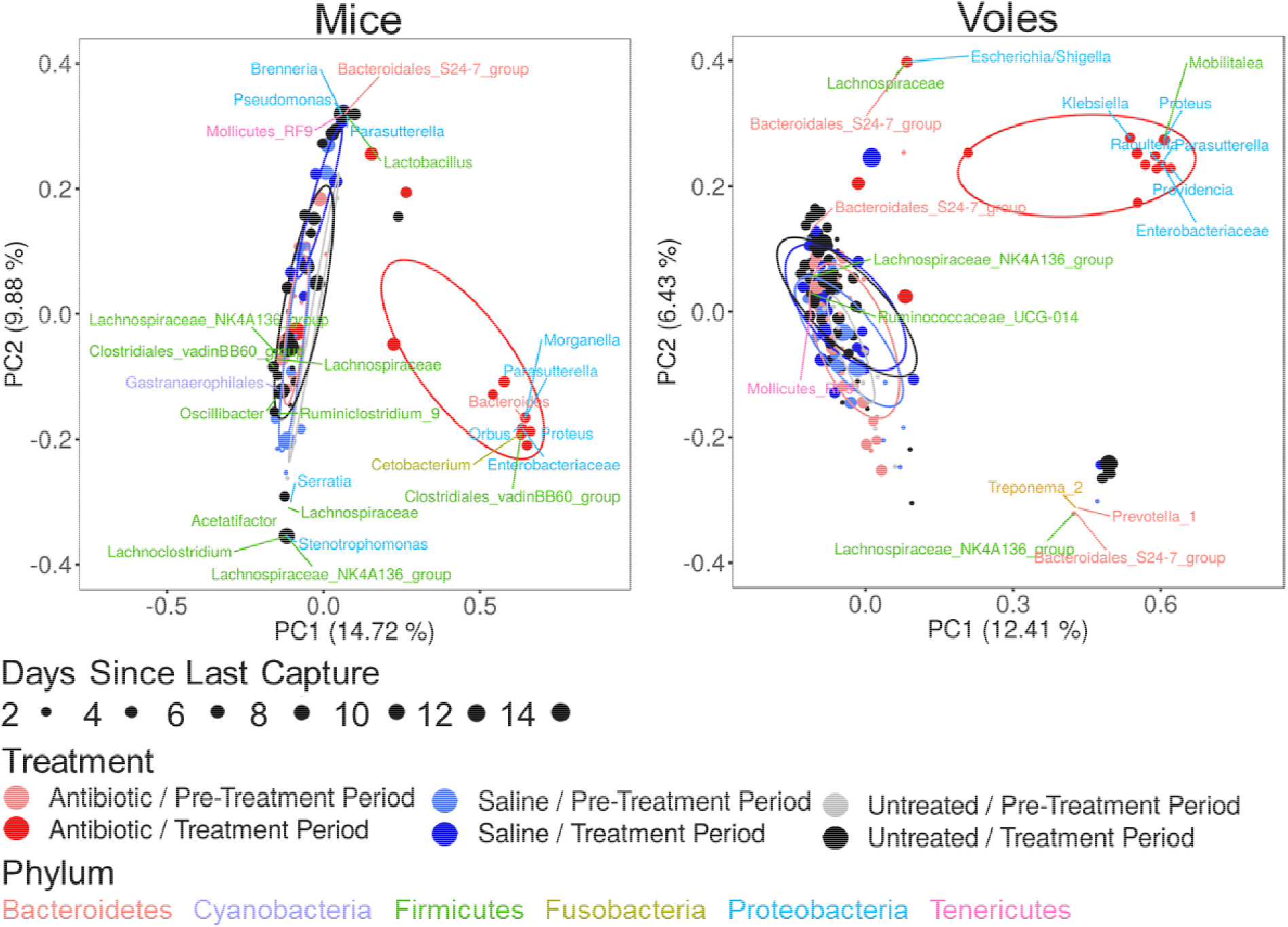
Effect of an antibiotic treatment on the composition of the gut microbiome. Principal coordinate analysis (PCoA) of Bray-Curtis distances among all samples collected from deer mice and red-backed voles. Treatment groups (Antibiotic, Saline, Untreated) before and during the treatment period are indicated by color and 95% confidence ellipses. Point size indicates the number of days since the last capture which, for the treatment period antibiotic group, represents the number of days since the last treatment. Phyla (color-coded) with the greatest weighted average scores in PCoA space are also plotted. Each point represents an individual sample.

**Fig.3.**
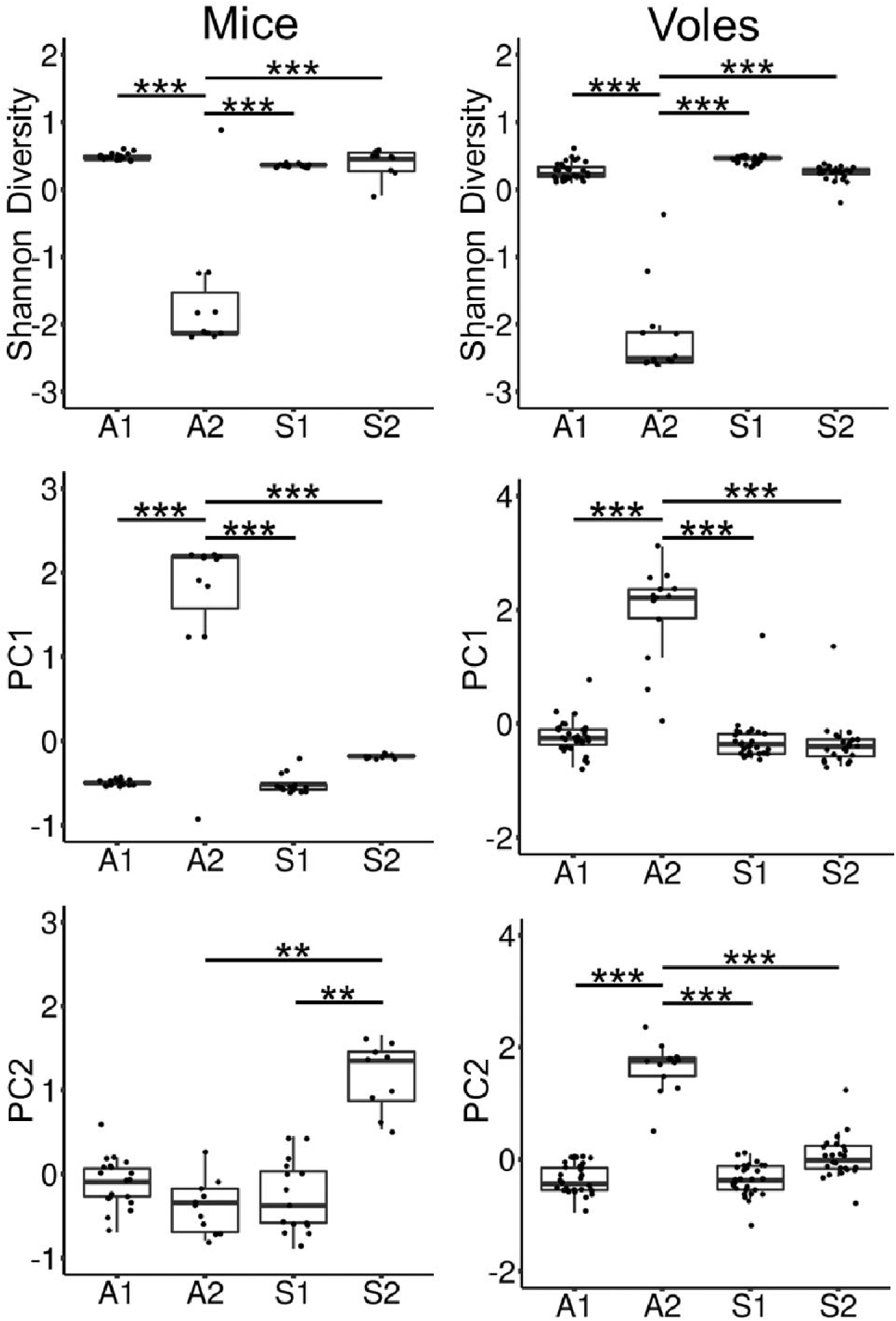
Effect of the antibiotic treatment on the composition of the gut microbiome of deer mice and red-backed voles given by the Shannon diversity and the scores of the first two axes from a PCoA of Bray-Curtis distances. Data are predicted values from separate mixed models for each species and between-group significance is assessed by pairwise comparisons of least squares means. A1=Antibiotic Pre-treatment, A2=Antibiotic Treatment, S1=Saline Pre-treatment, S2=Saline Treatment. ***P<0.001, **P<0.01.

In mice, weighted average scores of microbial taxa on ordination axes show that antibiotic-treated animals were mainly associated with an increase in relative abundance of taxa from the phylum Proteobacteria, with the greatest weight attributed to the genera *Morganella, Parasutterella, Orbus, Proteus*, and to other unknown members of the family Enterobacteriaceae (Fig. 2). This was met with a relative decrease in the phyla Firmicutes and Bacteroidetes. Antibiotic-treated animals also clustered with *Cetobacterium* of the phylum Fusobacteria, and an unknown Clostridiales taxon of the phylum Firmicutes. The increase in saline group PC2 was largely due to a relative decrease in Firmicutes, and a relative increase in Bacteroidetes, of which the order Bacteroidales had the greatest weight. This change was also associated with higher relative levels of the genera *Brenneria, Pseudomonas* and *Parasutterella* of the phylum Proteobacteria, and of the class Mollicutes of the phylum Tenericutes (Fig. 2). Unlike mice, the microbiome composition of voles did not change independently of the antibiotic treatment. The treatment resulted in an increase in the relative abundance of Proteobacteria, with the greatest weight attributed to the genera *Klebsiella, Parasutterella, Raoultella, Providencia, Proteus* of the family Enterobacteriaceae, and the genus *Mobilitalea* of the phylum Firmicutes (Fig. 2). The above results were largely supported by a differential expression analysis (Figs. S2, S3). Mice treated with the antibiotic showed a drastic loss in most taxa of the phyla Bacteroidetes and Firmicutes, with an increase in relative abundance of *Klebsiella sp., Parasutterella, Morganella morganii*, other Enterobacteriaceae sp. (Proteobacteria), *Fusobacterium ulcerans* (Fusobacteria), *Bacteroides sp*. and Bacteroidales (Bacteroidetes), and *Erysipelatoclostridium ramosum* (Firmicutes). The change in composition of the saline group from the pre-treatment to treatment period resulted from a decrease in Firmicutes which were mostly *Ruminoclostridium* (Clostridiales), and an increase in Bacteroidetes, mainly comprised of Bacteroidales (Fig. S2). For voles, taxa differentially expressed due to the treatment were mainly Proteobacteria and included *Escherichia/Shigella sp., Klebsiella sp., M. morganii, Raoultella sp., Proteus sp*., other unknown Enterobacteriaceae and *Enterobacter*, and *Parasutterella*. The treatment also caused an increase in an unknown *Clostridium* taxon, *Enterococcus sp., Erysipelotrichaceae*, and *Lactobacillus sp*. (Firmicutes; Fig. S3).

### Antibiotic Treatment Increases Exploration

Following the high vole mortality during the season (see next section), we restricted our analyses on the effects of antibiotics on behaviour to deer mice. We measured exploration (i.e. total distance travelled in the open field) and anxiety (i.e. the latency to enter the center of the open field; 29, 30) behaviour just before, and at the end of the treatment period. We found a significant effect of period on distance travelled in the open field in the antibiotic treatment group but no significant effect on anxiety, in deer mice (see Table S3). We thus conducted a path analysis to determine the most statistically supported causal path from the antibiotic treatment to the change in exploration. Two equivalent models best explained variation in distance travelled in the open field (Fig. 4). In both models, distance travelled decreased with period and increased with PC2, which was unrelated to the antibiotic treatment. However, in the first model, distance travelled increased (though marginally significant) with antibiotic (i.e. independent of its effect on the microbiome). In the second model, the increase in distance travelled resulted from the change in microbiome by the treatment.

**Fig.4.**
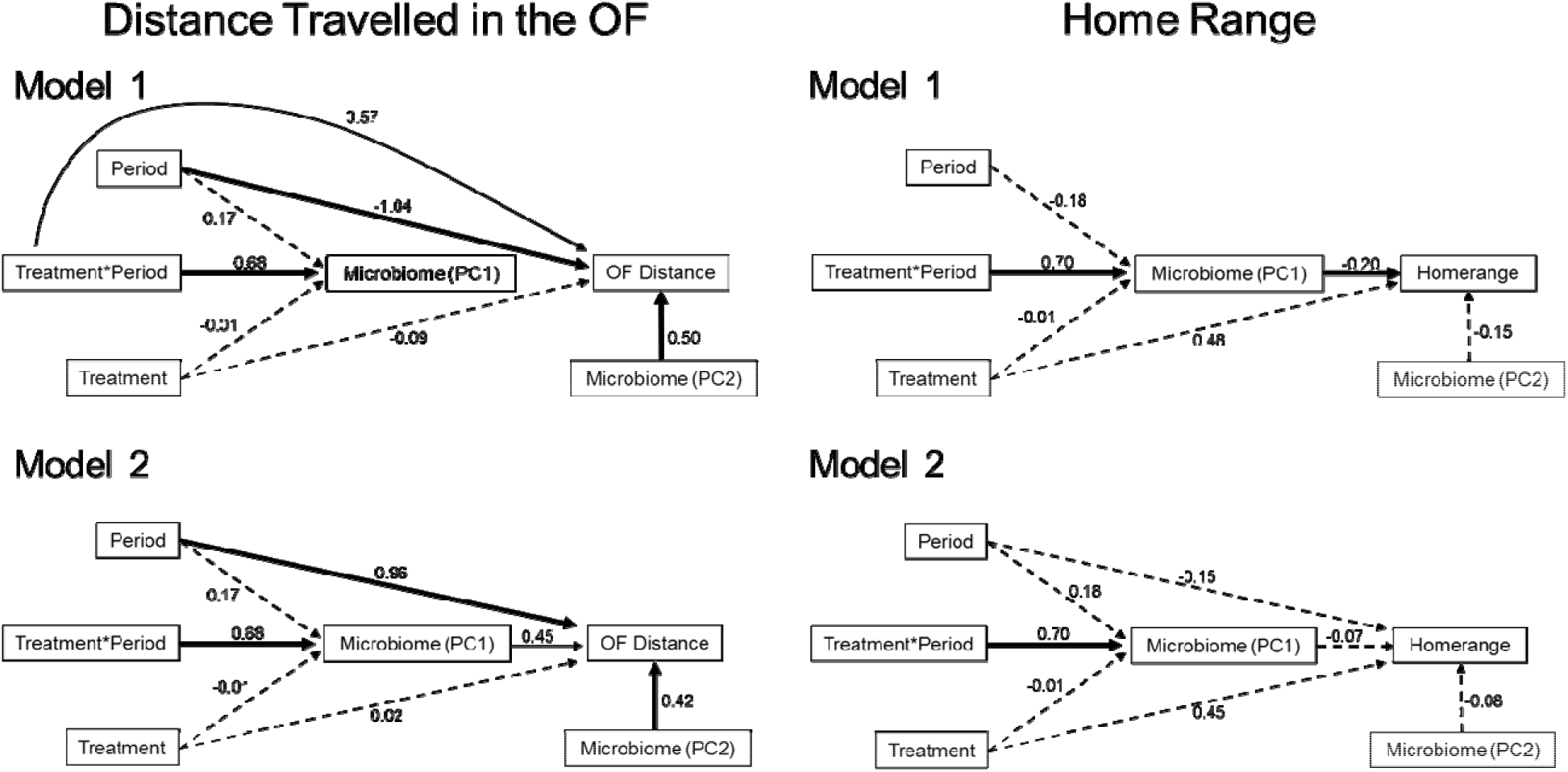
Path-analysis of the most likely causal relationships among treatment group (Antibiotic, Saline), period (Pre-Treatment, Treatment), microbiome (Scores PC1 and PC2 from PCoA of Bray-Curtis distances), and behaviour (OF distance and home range size) in deer mice. Two statistically equivalent models for each behaviour (model 1 and model 2) were retained after comparison of 13 hypothesized models. Values are path coefficients and significant paths are solid lines (bold = P<0.05, unbolded = P<0.1).

### Antibiotic Treatment Reduces Home Range Size

We measured the 95% kernel home range of each mouse during the pre- and post-treatment periods. We found significant effects of the antibiotic treatment on home-range size in deer mice (see Table S3). Two equivalent path analysis models best explained variation in home range size (Fig. 4). The first model included a reduction in home-range size resulting from the antibiotic-induced change in microbiome (Fig. 4). In the second model, the treatment modified the microbiome, but not home-range size. Given that the relationship between home range size and the microbiome (i.e. PC1) disappeared once we accounted for period (Fig. 4), this relationship was likely driven by the antibiotic treatment.

### Antibiotic Treatment Reduces Vole Survival

We compared survival during the treatment period of antibiotic-treated, saline-treated, and untreated animals with cox proportional hazards analysis (Therneau, 2015). Fifteen voles and no mice died in captivity. Five (1 Antibiotic, 4 Untreated) were found dead in the trap when traps were collected, and 10 (5 Antibiotic, 1 Saline, 4 Untreated) died during the manipulation or before release. The average (± SD) proportion of days animals were detected between the first and last detection during the treatment period was 81.5 ± 20.7% for mice, and 84.6 ± 19.0% for voles. We found no differential change in weight between antibiotic- and saline-treated mice or voles but mice increased in weight (t=2.7, p=0.02) and voles maintained a stable weight (t=-2.0, p=0.07) throughout the study (Fig. S7). For mice the antibiotic treatment did not change the hazard ratio when compared to the saline group (z=-0.44, *P*=0.66) or the untreated group (z=0.20, *P*=0.84, Fig. 5). Conversely, for voles, the antibiotic group showed a reduced hazard ratio compared to the untreated group (z=-3.1, *P*=0.002) and a marginally reduced hazard ratio compared to the saline group (z=-1.8, *P*=0.08, Fig. 5).

**Fig.5.**
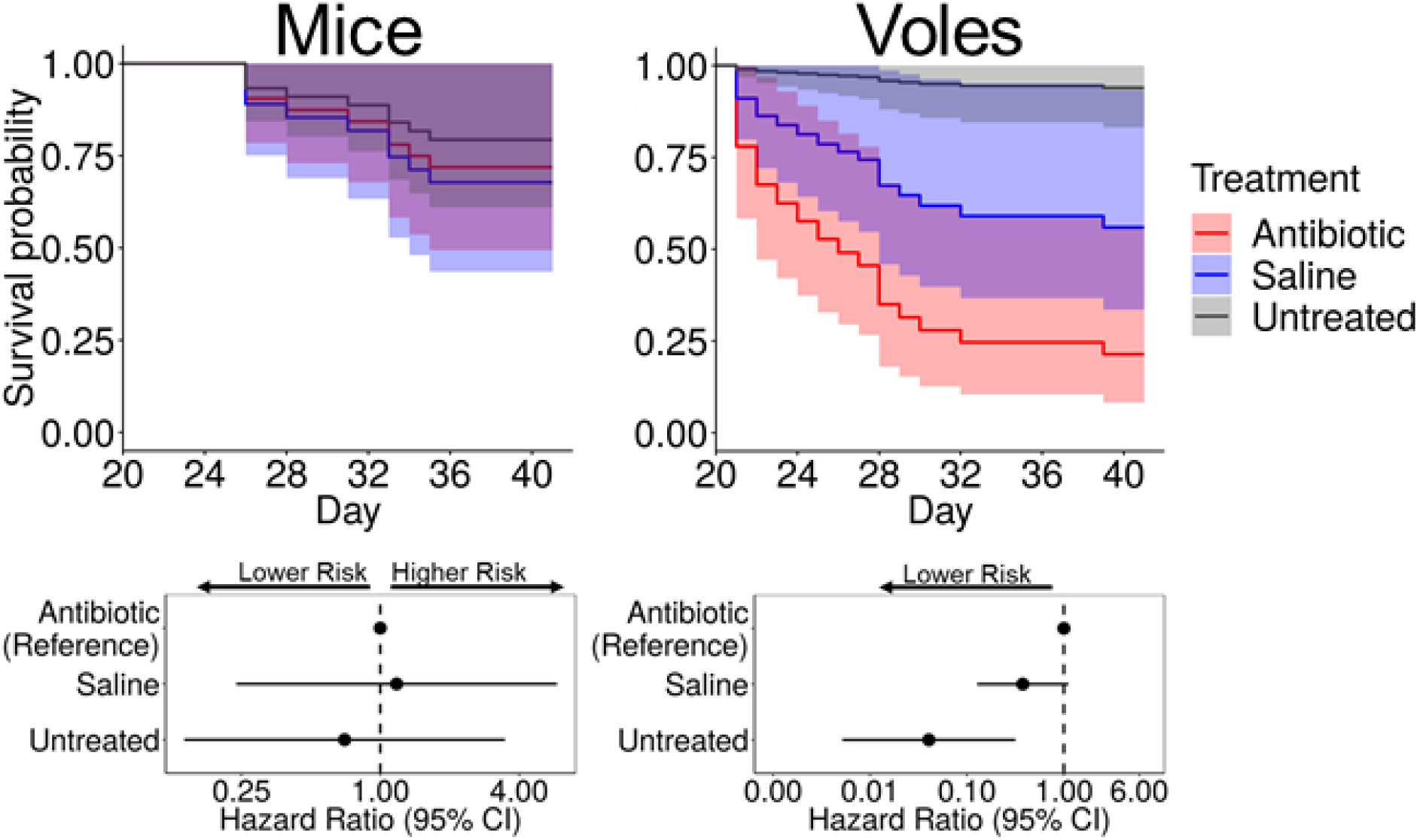
Comparison of survival of three treatment groups (Antibiotic, Saline, Untreated) during the treatment period using a Cox proportional hazards ratio analysis for deer mice and red-backed voles. Hazard ratios below 1.00 represent a lower likelihood of death, while values above 1.00 indicate a higher likelihood of death.

## DISCUSSION

Experimental studies have not yet addressed the potential impact of microbiome-mediated behavioural changes on the ecology and evolution of wild animals. Several ecological factors can influence the gut microbiome and behaviour of wild animals (Amato, 2013) and potentially overwhelm the effect of the microbiome on behaviour. Therefore, the goals of this study were to first, determine whether modifying the gut microbiome of wild mice and voles changes exploration and anxiety-related behaviours, and second, whether these changes have repercussions on the life history of the animals as measured by home-range size and survival.

### Antibiotic Treatment Reduces Microbiome Diversity

The antibiotic treatment reduced the diversity of the gut microbiome of both mice and voles. The gut microbiota of mice decreased in relative abundance of Bacteroides and Firmicutes and increased in relative abundance of Proteobacteria, namely *Klebsiella, Parasutterella, Morganella*, and other unknown Enterobacteriaceae. These same taxa also characterized the microbiome of antibiotic-treated voles, with the addition of *Escherichia/Shigella*. Interestingly, in mice, we also observed changes in the microbiome composition of the saline group independent of those in the treatment group, namely an increase in the relative abundance of Bacteroidetes, specifically Bacteroidales, and a decrease in that of Firmicutes, specifically Clostridiales.

### Antibiotic Treatment Increases Exploration

Exploration decreased in both antibiotic and saline-treated mice. This overall decrease in exploration in both treatment groups may have stemmed from stress induced by their daily manipulation. Chronic gavage has been shown to increase anxiety levels in laboratory mice (Gacias et al., 2016). In our study, anxiety (i.e. latency to enter the center) and exploration (i.e. distance travelled) were negatively correlated, suggesting that anxiety could have manifested as a decrease in exploration (Carter et al., 2013). However, this decrease was countered by a marginally significant increase in exploration only in antibiotic-treated mice (Fig. 4). This, at least in part, supports our hypothesis that the antibiotic-induced change in microbiome changed exploration-related behaviour. The ability of the microbiome to modulate exploration might help promote cohesion among animals that live in dense populations and new migrants entering a population. Transfer of bacteria among animals that live in groups promotes homogenization of microbiota which could favor homogenization of behaviours to promote positive interspecific interactions/cooperation. Other factors that change the microbiome may also change behaviour. The strong relationship between diet and the microbiome in many species means that seasonal changes in diet in wild animals could lead to microbiome-induced seasonal changes in behaviours that could potentially improve foraging success.

We also observed correlations between the gut microbiome and exploration that were unrelated to the antibiotic treatment. Specifically, the overall decrease in exploration with period was met with a decrease in the relative abundance of Firmicutes (Clostridiales) and an increase in the relative abundance of Bacteroidetes (Bacteroidales) (as seen in saline-treated mice) suggesting that either of these orders or both may be related to exploration behaviour. This is further supported by an overall positive correlation between exploration and PC2 which primarily represented a Firmicutes/Bacteroidetes gradient. Indeed, changes in the relative abundance of bacteria from the phylum Firmicutes have been linked to changes in social avoidance and exploration-related behaviours (Bercik et al., 2011; Gacias et al., 2016; Tochitani et al., 2016).

### Antibiotic Treatment Reduces Home Range Size

The antibiotic-induced change in microbiome resulted in a decrease in home range size. Additionally, unlike our results for open-field exploration, home range size did not correlate with the Firmicutes/Bacteroidetes gradient (i.e. PC2). This suggests that the increase in relative abundance of the Enterobacteriaceae was directly responsible for the change in home range. The Enterobacteriaceae, which include *Escherichia, Klebsiella* and *Morganella*, were the most expressed taxa in antibiotic-treated mice and voles. Their presence is associated with major depression in humans (Maes et al., 2008), which could potentially impact large-scale movement in wild animals if the same association is found in other species. They carry lipopolysaccharides (LPS) capable of inducing systemic and neuroinflammation (which can affect locomotor activity) and gut leakiness (Maes et al., 2008). Gut hyperpermeability further favors translocation of microbial metabolites like phenols into the periphery allowing them to reach the brain and alter neuro-activation (Gacias et al., 2016; Mohamadkhani, 2018). *Klebsiella pneumonia* and *M. morganii* are known to produce phenols at elevated concentrations (Saito et al., 2018). Immune inflammatory effects of LPS may also alter social behaviour as seen in autism (Mohamadkhani, 2018). We did not measure social behaviour but, given that the abundant taxa in antibiotic-treated mice have the potential to induce neurophysiological states related to depression and social withdrawal, a change in these behaviours may translate to a change in home range.

Finally, the additional ability of these taxa to produce serotonin (O’Mahony et al., 2015) could also allow them to act on the serotonergic system and modulate behaviour.

While it is counterintuitive that antibiotic-treated mice increased their exploration behaviour but reduced their home range, the relationship between behaviours measured in an open field and large-scale movements can be species- and context-specific. Open field exploration does not predict home range size in bank voles and starlings (Minderman et al., 2010; Schirmer et al., 2019) but it seems to in red squirrels and chipmunks (Boon et al., 2019; Montiglio et al., 2012). If the treatment is countering gavage-induced anxiety and potentially, social withdrawal, then treated mice may be less likely to withdraw if chased from a territory/resource by conspecifics and may also be more likely to visit an already occupied resource. This would preclude the need to visit farther stations. Additionally, animals with higher exploration or activity levels may compensate for a higher energetic output by reducing the area over which they are active. Finally, that variation in the microbiome composition (Firmicutes/Bacteroidetes gradient) unrelated to the antibiotic was also correlated with exploration but not with home range suggests that different bacterial taxa may influence different behaviours.

### Antibiotic Treatment Reduces Vole Survival

Survival of mice was unaffected by the antibiotic treatment and chronic gavage. In contrast, voles treated with antibiotics survived less than the control and the saline group. We could not differentiate survival from dispersal outside the capture grid, so the data suggest that the antibiotic did not affect emigration rates in mice. This is consistent with the related decrease in home range in this species. Although variation in exploration-related behaviour should influence individual predation risk and thus, survival (Smith and Blumstein, 2008; Wolf and Weissing, 2012), our continued presence on the study site may have deterred potential predators thereby removing the influence of susceptibility to predators on survival. This would most likely affect voles since mice are mainly active at night when we were absent, while voles are mainly diurnal. It is therefore surprising that the treatment lowered the survival of voles and not that of mice. This difference may indicate potential differences in host-microbiome interactions between the two species. Given that most of the voles that died during or just after the manipulation were antibiotic-treated and that the manipulation did not kill any mice, death not emigration is the likely cause of the disappearance in voles. Digestive malabsorption is not likely to have caused their lowered survival since they maintained a stable weight throughout the experiment. Voles may be more sensitive than mice to pathologies induced by the Enterobacteriaceae such as gastrointestinal low-grade inflammation or even septicemia. Additionally, while most species of *Escherichia* and *Shigella* are non-pathogenic, some are, and these genera were differentially abundant in voles but not in mice which may explain their differential survival. Finally, high population densities can lead to suppressed host immunity and fitness (Svensson et al., 2001). Voles had a relatively high population density while the density of mice was low and ten times lower than in the previous year indicating a recent crash in the population. Density-dependent effects may therefore have contributed to the lower sensitivity of mice and to the higher sensitivity of voles to both the gavage and the change in microbiome.

### Concluding Remarks

Antibiotics offer a practical and efficient way of manipulating the microbiome to understand its function. While widely used in laboratory studies, to our knowledge our study is the first to apply this method to wild populations. This study provides the first confirmation that the microbiome can directly modulate behaviour and aspects of life history in a population of wild animals. Administration of antibiotics reduced gut microbiome diversity of deer mice, favoring members of the Enterobacteriaceae. This change increased exploration weakly in opposition to a stronger likely stress-related decrease in exploration, and decreased home range, without impacting short-term survival. Aspects of the microbiome unrelated to the treatment were also correlated with exploration but not with home range in mice. The effect of the microbiome on behaviour may help drive behavioural adaptation in the face of changing environmental factors such as population structure and resource availability. Conversely, it may weaken populations if it affects ecologically relevant traits, or survival. Antibiotics also reduced gut microbiome diversity of red-backed voles, favoring once again the Enterobacteriaceae, with strong effects on vole survival, preventing us from testing the effect of the microbiome on vole behaviour. Our results suggest that host microbiome interactions depend on species and that different bacterial taxa may modulate or relate to different behaviours.

## MATERIALS AND METHODS

We conducted this study on Harbour Island in the Winnipeg river basin, Ontario (50°2.580’N, 94°40,436’W). The site consists of mixed boreal forest dominated by balsam fir (*Abies balsama*) paper birch (*Betula papyryfera*), white spruce (*Picea glauca*) bigtooth aspen (*Populus gradidentata*) and trembling aspen (*Populus tremuloides*).

We captured mice and voles with Longworth and BIOEcoSS (BIOEcoSS Ltd.) live traps, over a period of 41 days from 14 July – 23 August 2016. We placed 192 traps at 96 stations (8 x 12) distanced 10 m apart. The capture grid (9600 m^2^) was large enough to estimate individual home range sizes (Thompson et al., 2009; Vanderwel et al., 2010; Wolff, 1985; Wood et al., 2010). We conducted captures approximately every two days, weather permitting, opening the traps in the evening and checking and closing them at 6:00 am. Before each capture session we sterilized traps with 70% ethanol and baited them with exactly 4 g of carrot and 1 ml of peanut butter. After processing, we returned animals to the precise location they were captured. During a pre-treatment period (first 17 days) animals were captured but received no treatment. During the treatment period (day 18 to 41), animals were randomly assigned to either a group given saline, or a group given an antibiotic cocktail of Metronidazole (75 mg/kg), Ampicillin (100 mg/kg), Vancomycin (50 mg/kg), and Neomycin (50 mg/kg). This cocktail has been successfully used to deplete the gut microbiome and induce a GF phenotype in lab mice (Reikvam et al., 2011). We also included the antifungal Pimaricin (0.19 mg/kg) to stave off fungal infections (Bercik et al., 2011; Bravo et al., 2012; Foster and Neufeld, 2013). The duration and start/end dates of each period depended on each animal’s capture history.

We collected fecal samples from traps and collected two to three samples per animal throughout the study. Tools were systematically sterilized with 70% ethanol and flame just before sampling. Samples were placed in 99% ethanol, frozen at −20°C until the end of the study, then moved to a −80°C freezer. For each sample, we bead-beat 10 mg of fecal material in extraction kit ASL buffer (beat 1 min, rest 3 min, beat 1 min). We extracted DNA with the QIAGEN QIamp DNA Stool Minikit and randomly attributed samples to sequencing kit. The V4-V5 regions (~410 bp) of the 16S rRNA gene of extracted DNA were amplified with 515f/926r primers (Walters et al., 2015) and sequenced on an Illumina MiSeq at the Centre for Comparative Genomics and Evolutionary Bioinformatics (CGEB), Dalhousie University. We used the package DADA2 in R (R Core Team, 2018) to identify amplicon sequence variants (ASV) which were used as the fundamental unit of ecological analysis. It discriminates single nucleotide sequence variation by first modelling sequence error rate and correcting sequences for this error (Callahan et al., 2016). We used the SILVA database (V128) for taxonomic assignment using the RDP Naive Bayesian Classifier algorithm in DADA2. We removed ASVs with <100 sequences and rarefied ASVs to 4500 sequences per sample (Fig. S1). We first calculated Shannon’s diversity (alpha diversity) of each sample. We then Hellinger-transformed the ASV relative abundances and calculated the Bray-Curtis dissimilarity among the samples (Oksanen et al., 2019). We used principal coordinate analysis to plot pairwise dissimilarities in two dimensions and calculated the weighted average scores for taxa to identify those most associated with each PCoA axis (Oksanen et al., 2019). We overlaid taxa with the greatest weight onto PCoA plots. Rarefaction, diversity estimations and PCoAs were performed with package Vegan in R.

We measured anxiety and exploration behaviour with an open-field test just before, and at the end of the treatment period (Montiglio et al., 2010; Wolfer et al., 2004). The arena was cleaned with 70% ethanol before each trial. We could not measure post-treatment behaviour of animals that were not recaptured at the end of the study. Animals were individually placed in the same corner of an empty arena (46 cm^3^) and video-recorded from above for five minutes. We tracked their movement in the arena and extracted behavioural variables using Ethovision software (V.9.0, Noldus Information Technology, Wageningen, The Netherlands). We measured exploration as the total distance travelled in the open field. We also defined a central square in the arena (23 cm^2^) which is perceived by the animal as a riskier zone, and measured levels of anxiety as the latency to enter this zone.

To measure home range, when first captured, we outfitted each animal with a subcutaneous passive integrated transponder (PIT) tag. This allowed us to rapidly identify recaptured animals, and to follow their movement within the capture grid. We deployed 16 RFID antennae at equal distances throughout the capture grid from July 25 (Day 12) to August 23 (Day 41). The RFID system is described in the supplementary information section. When an animal passed through the antenna ring, a data logger recorded its identity and the time of detection. To measure home range, we counted multiple consecutive detections for an individual at the same RFID station and on the same day as a single detection. We calculated the 95% kernel home range for each individuals’ pre-treatment and treatment period with the package adehabitatHR in R (Calenge, 2006).

To assess the impact of the antibiotics on the gut microbiome of each species and determine which of the PCoA axes was associated with the treatment, we ran separate mixed models on alpha diversity and on the scores for each of the first two PCoA axes (PC1, PC2) for each species using package nlme in R. We started with the fully parameterized model which included treatment, period, days since last capture and their interactions as fixed effects, and ID as a random effect. We performed model selection by sequentially removing a term and using the log likelihood ratio to test the simplified model against the full model. We validated all final models with respect to model assumptions. We used DESEQ2 (package DESEQ2 in R) differential expression analysis to identify microbial taxa that showed the greatest treatment-induced change in abundance and their log2 fold change (Love et al., 2014). For each pairwise contrast in this analysis, the 30 most significant taxa were compiled, and we generated a heatmap of their log transformed normalized counts (see supplementary material). Factors included in the DESEQ2 model design were treatment, period, and their interaction.

Given the high mortality in voles throughout the season we restricted analyses of behaviour and home-range size to deer mice. We sought to identify the most likely causal model that explained the relationships between the variables. For this we used structural equation modelling (package PiecewiseSEM in R; 52). We first ran mixed models (package nlme in R) of open-field behaviour and home-range size as a function of period, PC1, PC2 and their interactions with period to determine which variables to include in the SEMs. We compared full and simplified mixed models using likelihood ratio tests. We then sought to identify the most likely causal model that explained the relationships between the variables. For this we used structural equation modelling (Lefcheck, 2016). We compared 13 potential causal models based on model Fisher’s C and AIC and obtained standard estimates for each pairwise relationship between the variables. For all models, we included ID as a random variable and, in models for home range, we included number of RFID detections as a weighting factor to correct for the potential effect of the number of detections on the accuracy of the home range estimation.

Finally, we measured survival during the treatment period (days 18-41) using Cox proportional hazards analysis with the package survival in R (Therneau, 2015). Our data better suited this modelling approach, as opposed to capture-mark-recapture models due to the high detection rate of both mice and voles (see results). For this analysis, we had enough data to include untreated animals. Most untreated mice were females, so we included all adult-untreated mice. There were enough untreated adult male voles that we excluded female and subadult male voles from the analysis. We pooled the capture and RFID detection data to obtain the first and last detections of each animal. We specified five days as the threshold after which an animal was considered either dead or no longer in the study site. We were confident in this assumption given the high detection rate of our animals. We excluded all animals that died while captive. We ran separate models for each species and included treatment, number of treatments, and their interaction as fixed effect in the models.

## ACKNOWLEDGMENTS

Animal care and experimental procedures were performed in accordance with protocols approved by the Comité institutionnel de protection des animaux (CIPA #917). We thank S. Zhao, C. Pelletier, R. Pedneault for their assistance and dedication in the field and K. and B. Hall for their logistical support in the field. C. Negre performed the DNA extractions and assisted with analyses. We thank A. Fontaine, D. Bourget, P. Bergeron for their suggestions and support with survival analyses. A. Mahamane provided support in selecting and obtaining the antibiotics. J Jameson received an NSERC Alexander Graham Bell Canada Graduate Scholarship, and S. Zhao and C. Pelletier received NSERC-USRA fellowship. This research was funded by Natural Sciences and Engineering Research Council of Canada (NSERC) Discovery grants (Réale and Kembel), a Canada Research Chair (Kembel), an American Society of Mammalogists Grant-in-Aid of Research (Jameson) and an Animal Behavior Society Student Research Grant (Jameson).

