## Supplemental material for "Gut microbiome modulates behaviour and life history in two wild rodents"

SUPPLEMENTARY MATERIAL

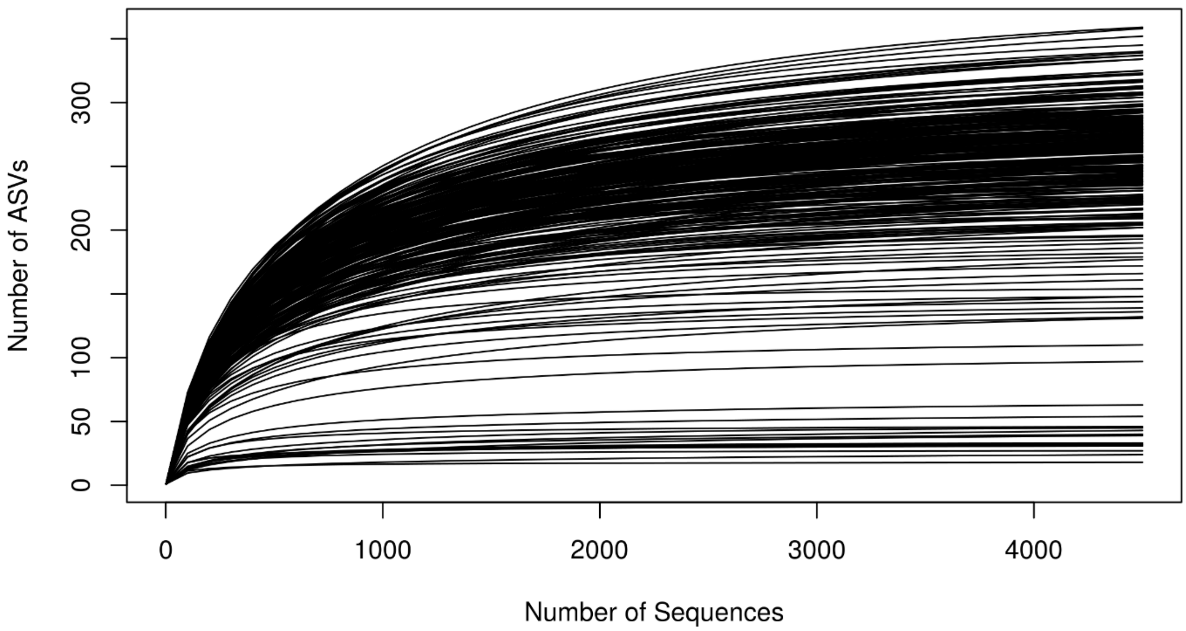


Figure S1. Rarefaction curves for all samples collected depicting the number of amplicon sequence variants (ASV) per number of sequences in each sample.

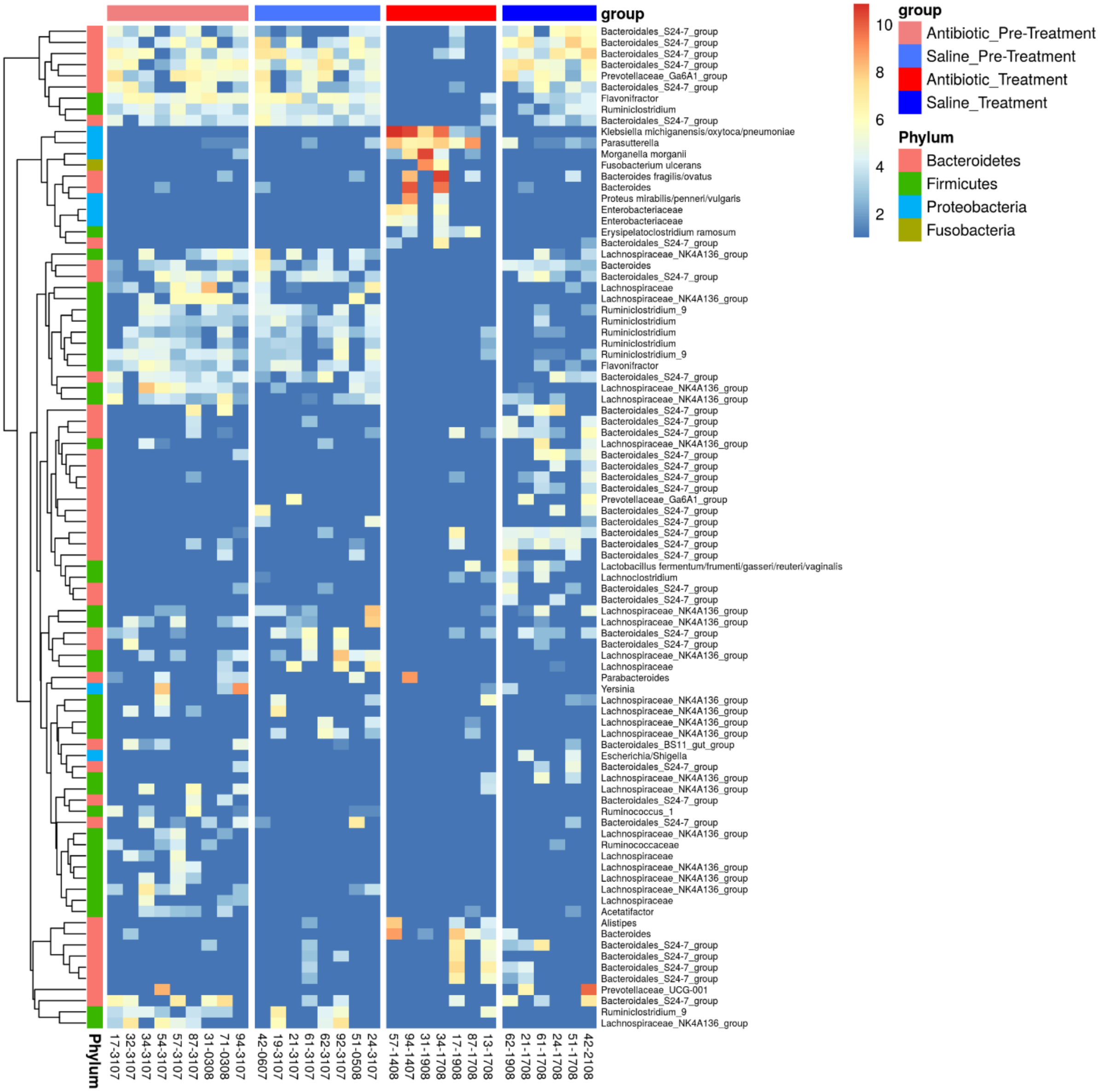


Figure S2. We used DESEQ2 differential expression analysis to identify microbial taxa that showed the greatest change in abundance among treatments and periods in deer mice (*Peromyscus maniculatis*). For each pairwise contrast in this analysis, the 30 most significant taxa (adjusted P-values < 0.05) were compiled, and we generated the above heatmap with their log transformed normalized counts.

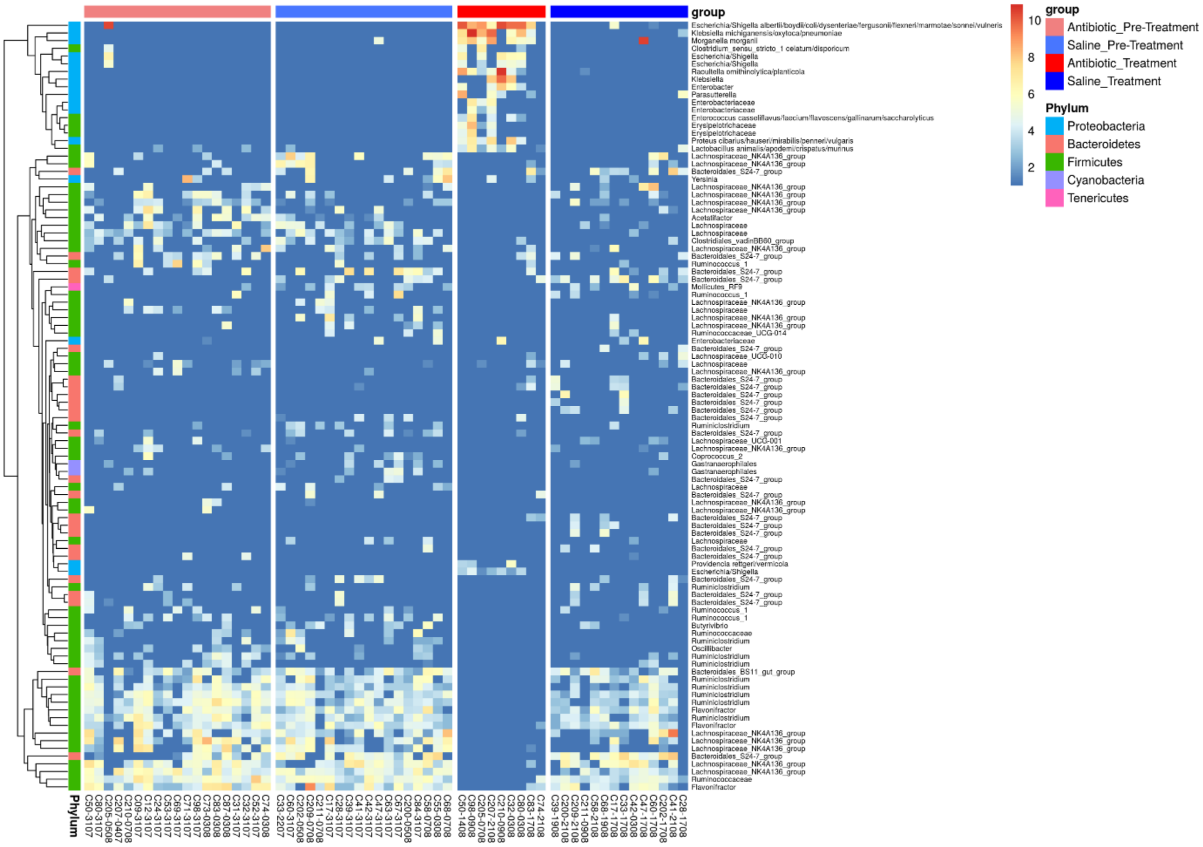


Figure S3. We used DESEQ2 differential expression analysis to identify microbial taxa that showed the greatest change in abundance among treatments and periods in red-backed voles (*Myodes gapperi*). For each pairwise contrast in this analysis, the 30 most significant taxa (adjusted P-values < 0.05) were compiled, and we generated the above heatmap with their log transformed normalized counts.

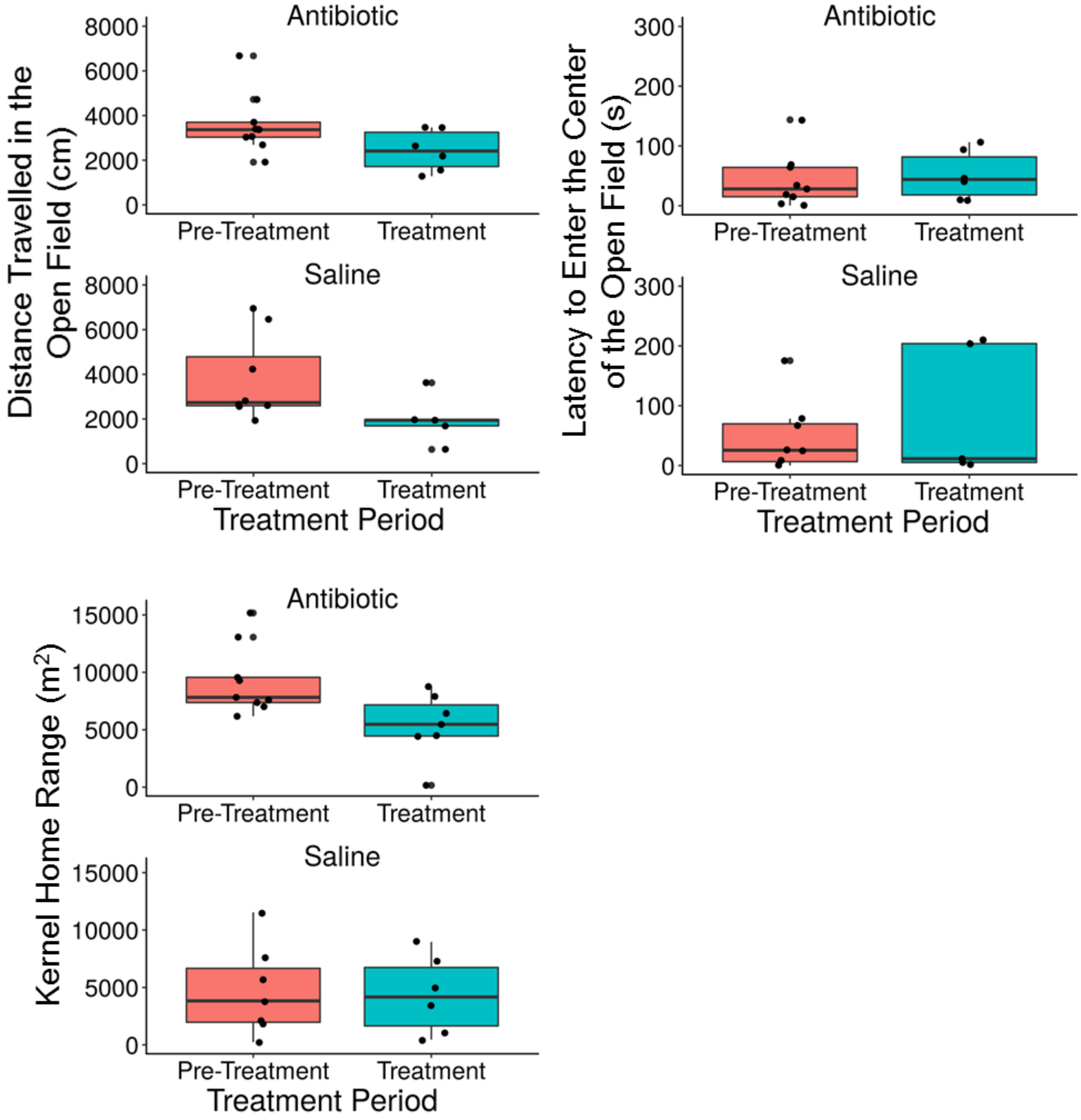


Figure S4. Deer mouse (*Peromyscus maniculatis*) behaviour data (distance travelled in the open field (cm), latency to enter the center of the open field (s), and kernel home range ($m^2$)) as a function of antibiotic treatment and sampling period.

Table S1. Effect of the antibiotic treatment  on gut microbiome Shannon diversity and composition (scores from the first two axes of a PCoA of Bray-Curtis distances) of deer mice (*Peromyscus maniculatus*) and red-backed boles (*Myodes gapperi*)

| | | Mice | | | | | Voles | | | | |
|---|---|---|---|---|---|---|---|---|---|---|---|
| | Effects | Value | SE | DF | t | p | Value | SE | DF | t | p |
| Shannon Diversity | (Intercept) | 0.51 | 0.10 | 30 | 5.01 | <0.001 | 0.32 | 0.09 | 56 | 3.52 | 0.001 |
| | treatment | -0.14 | 0.14 | 16 | -1.01 | 0.327 | 0.13 | 0.13 | 35 | 1.04 | 0.304 |
| | period | -2.48 | 0.16 | 30 | -15.50 | **<0.001** | -2.79 | 0.17 | 56 | -16.30 | **<0.001** |
| | days since the last capture | 0.06 | 0.13 | 30 | 0.44 | 0.663 | 0.17 | 0.12 | 56 | 1.42 | 0.162 |
| | treatment : period | 2.62 | 0.24 | 30 | 10.90 | **<0.001** | 2.64 | 0.22 | 56 | 11.99 | **<0.001** |
| | treatment : days since the last capture | -0.05 | 0.15 | 30 | -0.35 | 0.727 | -0.18 | 0.14 | 56 | -1.30 | 0.200 |
| | period : days since the last capture | 0.78 | 0.17 | 30 | 4.59 | **<0.001** | 0.97 | 0.26 | 56 | 3.76 | **<0.001** |
| | treatment : period : days since the last capture | -1.04 | 0.25 | 30 | -4.11 | **<0.001** | -1.06 | 0.28 | 56 | -3.73 | **<0.001** |
| PC1 | (Intercept) | -0.49 | 0.08 | 30 | -6.27 | 0.000 | -0.26 | 0.14 | 56 | -1.92 | 0.060 |
| | treatment | -0.01 | 0.11 | 16 | -0.09 | 0.932 | 0.00 | 0.19 | 35 | -0.01 | 0.996 |
| | period | 2.52 | 0.12 | 30 | 20.36 | **<0.001** | 2.46 | 0.19 | 56 | 13.12 | **<0.001** |
| | days since the last capture | 0.01 | 0.10 | 30 | 0.09 | 0.928 | -0.17 | 0.12 | 56 | -1.46 | 0.151 |
| | treatment : period | -2.20 | 0.19 | 30 | -11.88 | **<0.001** | -2.59 | 0.23 | 56 | -11.17 | **<0.001** |
| | treatment : days since the last capture | 0.07 | 0.12 | 30 | 0.63 | 0.535 | 0.16 | 0.14 | 56 | 1.15 | 0.254 |
| | period : days since the last capture | -0.87 | 0.13 | 30 | -6.60 | **<0.001** | -1.01 | 0.25 | 56 | -4.00 | **<0.001** |
| | treatment : period : days since the last capture | 0.80 | 0.20 | 30 | 4.08 | **<0.001** | 1.07 | 0.28 | 56 | 3.81 | **<0.001** |
| PC2 | (Intercept) | -0.01 | 0.22 | 33 | -0.06 | 0.955 | -0.35 | 0.16 | 56 | -2.19 | 0.033 |
| | treatment | -0.25 | 0.33 | 16 | -0.76 | 0.456 | -0.06 | 0.22 | 35 | -0.28 | 0.782 |
| | period | -0.54 | 0.31 | 33 | -1.77 | 0.085 | 2.21 | 0.27 | 56 | 8.29 | **<0.001** |
| | days since the last capture | 0.24 | 0.11 | 33 | 2.08 | **0.046** | 0.01 | 0.18 | 56 | 0.08 | 0.935 |
| | treatment : period | 1.79 | 0.44 | 33 | 4.09 | **<0.001** | -1.80 | 0.33 | 56 | -5.38 | **<0.001** |
| | treatment : days since the last capture | - | - | - | - | - | -0.02 | 0.20 | 56 | -0.08 | 0.933 |
| | period : days since the last capture | - | - | - | - | - | -0.65 | 0.37 | 56 | -1.72 | **0.091** |
| | treatment : period : days since the last capture | - | - | - | - | - | 0.90 | 0.42 | 56 | 2.17 | **0.035** |

Table S2. Pairwise comparisons testing the interaction between the treatment (antibiotic or saline) and period (pre-treatment or treatment) on gut microbiome Shannon diversity and composition (scores from the first two axes of a PCoA of Bray-Curtis distances) of deer mice (*Peromyscus maniculatus*). A:antibiotic, S: saline, 1: pre-treatment period, 2: treatment period.

| | | Mice | | | | | Voles | | | | |
|---|---|---|---|---|---|---|---|---|---|---|---|
| | contrast | Estimate | SE | DF | t | p | Estimate | SE | DF | t | p |
| Shannon Diversity | **A1 vs A2** | 2.48 | 0.16 | 30 | 15.50 | **<0.001** | 2.80 | 0.17 | 56 | 16.30 | **<0.001** |
| | A1 vs S1 | 0.14 | 0.14 | 16 | 1.01 | 0.745 | -0.13 | 0.13 | 35 | -1.04 | 0.725 |
| | A1 vs S2 | 0.01 | 0.18 | 16 | 0.07 | 1.000 | 0.02 | 0.14 | 35 | 0.16 | 0.999 |
| | **A2 vs S1** | -2.34 | 0.16 | 16 | -14.70 | **<0.001** | -2.93 | 0.17 | 35 | -16.97 | **<0.001** |
| | **A2 vs S2** | -2.47 | 0.19 | 16 | -12.78 | **<0.001** | -2.77 | 0.18 | 35 | -15.15 | **<0.001** |
| | S1 vs S2 | -0.13 | 0.18 | 30 | -0.74 | 0.882 | 0.16 | 0.14 | 56 | 1.14 | 0.666 |
| PC1 | A1 vs S1 | 0.01 | 0.11 | 16 | 0.09 | 1.000 | 0.00 | 0.19 | 35 | 0.01 | 1.000 |
| | **A1 vs A2** | -2.52 | 0.12 | 30 | -20.36 | **<0.001** | -2.46 | 0.19 | 56 | -13.12 | **<0.001** |
| | A1 vs S2 | -0.31 | 0.14 | 16 | -2.22 | 0.159 | 0.13 | 0.21 | 35 | 0.64 | 0.920 |
| | **S1 vs A2** | -2.53 | 0.12 | 16 | -20.56 | **<0.001** | -2.46 | 0.23 | 35 | -10.54 | **<0.001** |
| | S1 vs S2 | -0.32 | 0.14 | 30 | -2.30 | 0.120 | 0.13 | 0.14 | 56 | 0.95 | 0.779 |
| | **A2 vs S2** | 2.21 | 0.15 | 16 | 14.81 | **<0.001** | 2.59 | 0.24 | 35 | 10.65 | **<0.001** |
| PC2 | A1 vs S1 | 0.25 | 0.33 | 16 | 0.77 | 0.869 | 0.06 | 0.23 | 35 | 0.28 | 0.992 |
| | **A1 vs A2** | 0.54 | 0.31 | 33 | 1.77 | 0.304 | -2.21 | 0.27 | 56 | -8.29 | **<0.001** |
| | A1 vs S2 | -1.00 | 0.38 | 16 | -2.64 | 0.076 | -0.35 | 0.25 | 35 | -1.42 | 0.497 |
| | **S1 vs A2** | 0.29 | 0.37 | 16 | 0.80 | 0.854 | -2.27 | 0.29 | 35 | -7.84 | **<0.001** |
| | S1 vs S2 | -1.25 | 0.33 | 33 | -3.80 | **0.003** | -0.41 | 0.20 | 56 | -2.03 | 0.191 |
| | **A2 vs S2** | -1.54 | 0.40 | 16 | -3.83 | **0.007** | 1.86 | 0.31 | 35 | 6.10 | **<0.001** |

Table S3. Effect of the antibiotic-mediated change in gut microbiome on open-field behaviours and home-range size in deer mice (*Peromyscus maniculatus*). Results from mixed models testing for an effect of the gut microbiome composition given by the scores of the first two axes from a PCoA of Bray-Curtis distances on behaviours measured in an open field and on the 95% kernel home range. We ran separate models for each treatment group (Antibiotic or Saline). Full models were *Behaviour ~ period + PC1 + PC2 + PC1:period* for models on open-field behaviours and *Behaviour ~ period + PC1 + PC2 + PC1:period + PC2:period* for models on home range. We present here the results for the simplified models. Significant (≤0.05) p-values are bolded.

| Group | Term | Estimate | SE | DF | t-value | p-value |
|---|---|---|---|---|---|---|
| Antibiotic | **Distance Travelled ~ Period + PC1 + PC2** | | | | | |
| | | Estimate | SE | DF | t-value | p-value |
| | Intercept | 0.677 | 0.336 | 9 | 2.015 | 0.075 |
| | Period (Pre-Treatment) | -1.893 | 0.458 | 2 | -4.130 | **0.054** |
| | PC1 | 0.547 | 0.242 | 2 | 2.262 | 0.152 |
| | PC2 | 0.487 | 0.202 | 2 | 2.407 | 0.138 |
| Saline | **Distance Travelled ~ Period + PC1 + PC2 + Period*PC1** | | | | | |
| | Intercept | 0.461 | 0.292 | 7 | 1.579 | 0.158 |
| | Period (Pre-Treatment) | -1.861 | 0.505 | 1 | -3.683 | 0.169 |
| | PC1 | -1.180 | 0.581 | 1 | -2.032 | 0.291 |
| | PC2 | 1.155 | 0.440 | 1 | 2.622 | 0.232 |
| | Period*PC1 | 0.984 | 0.526 | 1 | 1.872 | 0.312 |
| Antibiotic | **Latency to Enter Center ~ Period + PC1 + PC2** | | | | | |
| | | Estimate | SE | DF | t-value | p-value |
| | Intercept | -0.016 | 0.385 | 9 | -0.042 | 0.968 |
| | Period (Pre-Treatment) | 0.006 | 0.439 | 2 | 0.015 | 0.990 |
| | PC1 | 0.229 | 0.233 | 2 | 0.983 | 0.429 |
| | PC2 | -0.054 | 0.201 | 2 | -0.266 | 0.815 |
| Saline | **Latency to Enter Center ~ Period + PC1 + PC2** | | | | | |
| | Intercept | 0.068 | 0.411 | 7 | 0.165 | 0.874 |
| | Period (Pre-Treatment) | -0.176 | 0.754 | 2 | -0.234 | 0.837 |
| | PC1 | 0.705 | 0.592 | 2 | 1.190 | 0.356 |
| | PC2 | -0.346 | 0.619 | 2 | -0.560 | 0.632 |
| Antibiotic | **Home Range ~ Period + PC1 + PC2 + Period*PC2** | | | | | |
| | | Estimate | SE | DF | t-value | p-value |
| | Intercept | 1.386 | 0.309 | 9 | 4.492 | 0.002 |
| | Period (Pre-Treatment) | -2.934 | 0.511 | 2 | -5.742 | **0.029** |
| | PC1 | 1.230 | 0.287 | 2 | 4.280 | **0.051** |
| | PC2 | -0.756 | 0.169 | 2 | -4.467 | **0.047** |
| | Period*PC2 | 1.045 | 0.272 | 2 | 3.842 | 0.062 |
| Saline | **Home Range ~ Period + PC1** | | | | | |
| | Intercept | 0.035 | 0.430 | 6 | 0.081 | 0.938 |
| | Period (Pre-Treatment) | -0.262 | 0.120 | 4 | -2.178 | 0.095 |
| | PC1 | -0.071 | 0.102 | 4 | -0.702 | 0.521 |

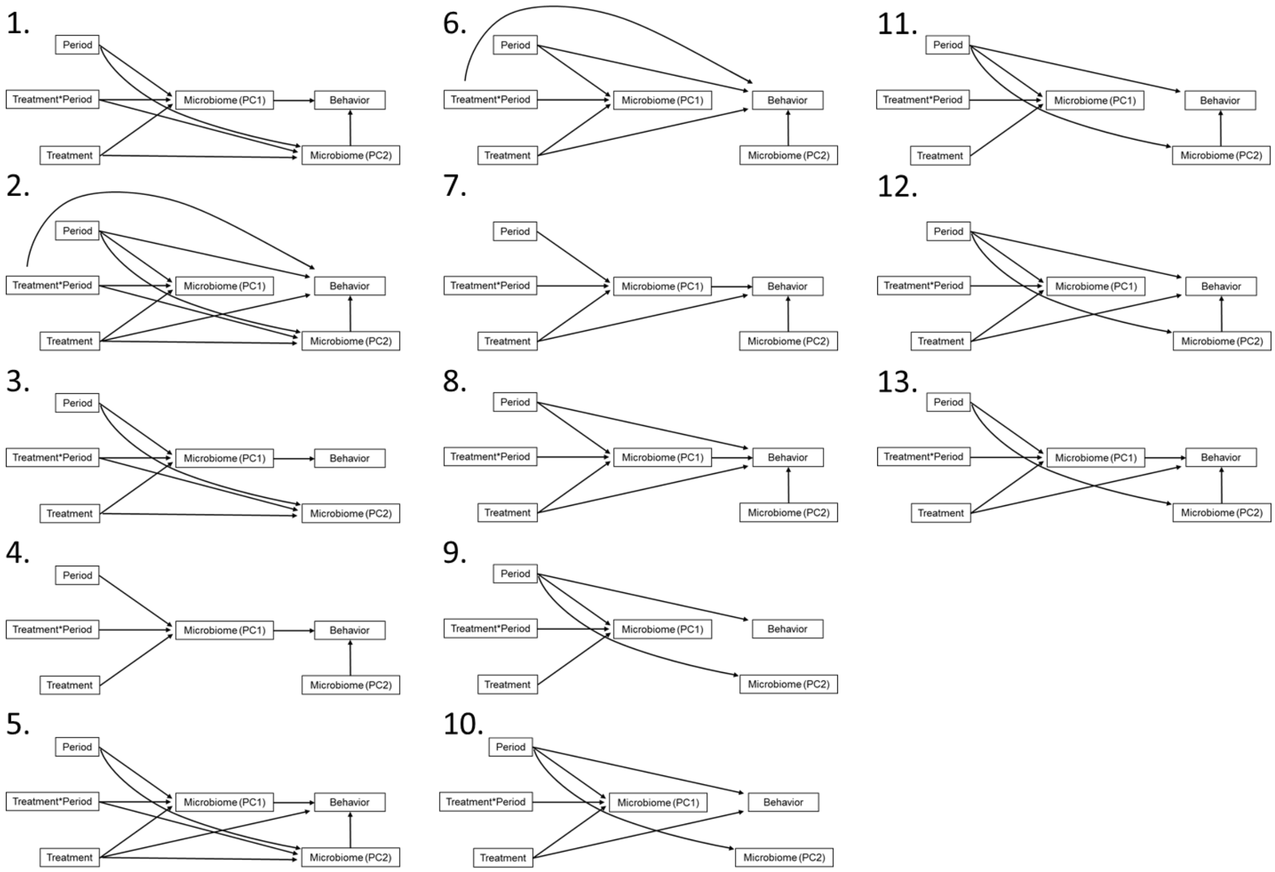


Figure S5. Path analysis models tested.

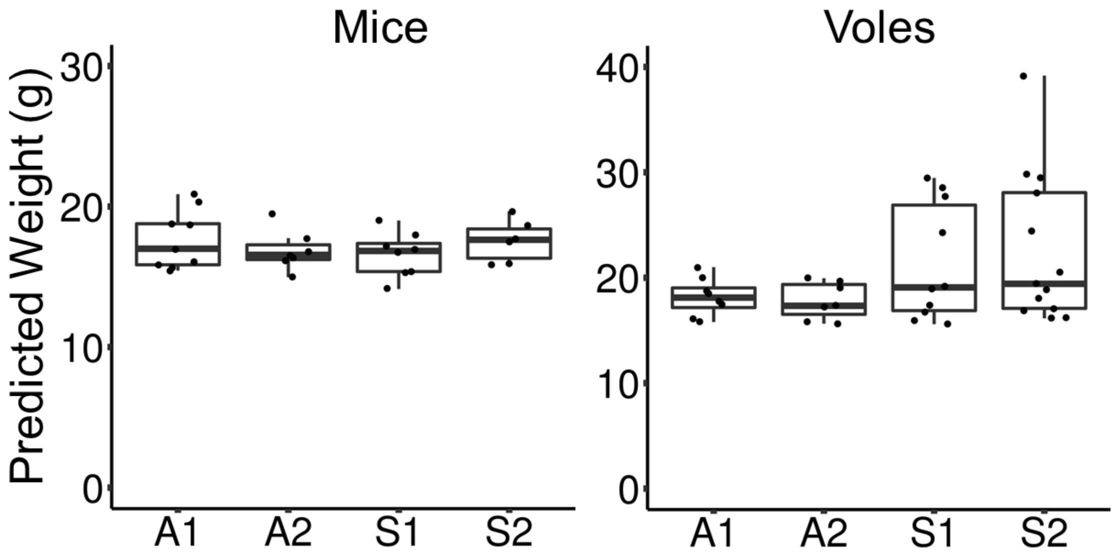


Figure S7. Predicted values for weight from mixed models testing for an effect of the antibiotic treatment on weight. There was no effect of the antibiotic on weight. Mice increased in weight from the pre-treatment to treatment period. The weight of voles remained stable throughout the experiment.

Description of RFID dataloggers

Dataloggers consisted of a ring-shaped antenna placed onto a wooden arena (20 x 20 x 5 cm). The arena contained closed compartments filled with peanut butter. The compartments were perforated so animals could smell the bait but not eat it. We regularly replaced the bait in these compartments. The floor of the arena was a plastic grid elevated 1-2 cm above the ground to prevent accumulation of feces within the arena and prevent unwanted cross-contamination among animals.
